# Data-efficient protein mutational effect prediction with weak supervision by molecular simulation and protein language models

**DOI:** 10.1101/2025.04.08.647800

**Authors:** Teppei Deguchi, Nur Syatila Ab Ghani, Yoichi Kurumida, Shinji Iida, Kaito Kobayashi, Yutaka Saito

## Abstract

Machine learning-based protein mutational effect prediction is widely used in protein engineering and pathogenicity prediction, but training data scarcity remains a major challenge due to high costs of experimental measurements. A previous study proposed data augmentation using computational estimates by molecular simulation. However, this approach has been limited to predicting mutational effects on thermostability. Here, we present a new data augmentation method that combines molecular simulation with zero-shot prediction computed by protein language models. These computational estimates serve as “weak” training data to supplement experimental training data. Our method dynamically adjusts the weight and inclusion of weak training data based on available experimental training data. This reduces potential negative impacts of weak training data while extending applicability to diverse protein properties such as binding affinity and enzymatic activity. Benchmark tests demonstrate that our method improves prediction accuracy particularly when experimental training data are scarce. These results indicate the capability of our approach to advance protein engineering and pathogenicity prediction in small data regimes.

## Introduction

Machine learning (ML)-based protein mutational effect prediction has been successfully applied to protein engineering and pathogenicity prediction(1-6). The task takes a mutant sequence as input and predicts its mutational effect, defined as the change of protein property of interest such as thermostability, binding affinity, and enzymatic activity compared to the wild type. ML models are trained using mutant sequences and their experimentally measured mutational effects(7-9).

To make accurate ML models, a large amount of training data is desired while it is often unavailable due to high costs of experimental measurements(8, 9). Therefore, data-efficient approaches are necessary to achieve accurate prediction with limited training data.

Augmentation using computationally generated data is an efficient way to virtually expand the amount of training data. In a previous study(10), molecular simulation was used as a means of data augmentation for thermostability prediction. Their method, mGPfusion, uses Rosetta molecular simulation software (11) to compute thermostability changes for all possible single-residue mutants. These computational estimates are integrated with experimentally measured thermostability to expand training data for ML models, thereby improving prediction accuracy. This method is regarded as data augmentation using computational estimates by molecular simulation, which could be a versatile approach for data-efficient mutational effect prediction. However, the previous study (10) was limited to thermostability prediction, and applications to other protein properties have not been explored.

More recently, protein language models (pLM) have been used to estimate the mutational effect based on the likelihood ratio of mutant and wild-type sequences(12, 13). This method is called zero-shot prediction since it works without using experimentally measured training data. Mutational effects estimated by zero-shot prediction are reported to correlate with diverse protein properties (13) although they have not yet been applied for data augmentation.

Here, we present a new data augmentation method that combines molecular simulation and zero-shot prediction using pLM. These computational estimates serve as “weak” training data to supplement experimental training data. Unlike the previous study (10) limited to thermostability, we aim to improve the prediction accuracy of mutational effects on diverse protein properties. Using Rosetta and ESM-2 (12) as pLM, we design a hybrid score that combines both estimates to enhance accuracy and robustness.

A key challenge in using weak training data is that inaccurate computational estimates can sometimes degrade ML model performance. To mitigate this issue, we develop algorithms that dynamically adjust the weight of weak training data and decide whether or not to include them based on available experimental training data.

Our method successfully improves the prediction accuracy on various proteins and their properties including protein abundance, binding affinity and enzymatic activity, surpassing ML models trained sorely with experimental data. The accuracy improvement is especially remarkable when experimental training data are scarce. Furthermore, we demonstrate that our method is capable of predicting the effects of double-residue mutations using single-residue mutants as training data.

## Results

### Methods overview

In this study, we developed a new method for protein mutational effect prediction with data augmentation using computational estimates by molecular simulation and pLM’s zero-shot prediction (Figure 1a). We employed Rosetta for molecular simulation and ESM-2 as pLM. Mutational effects estimated by Rosetta and ESM-2 are regarded as weak training data integrated with experimental training data. Using these integrated training data, the ML model is constructed to predict the mutational effect from a mutant sequence.

**Figure 1.**
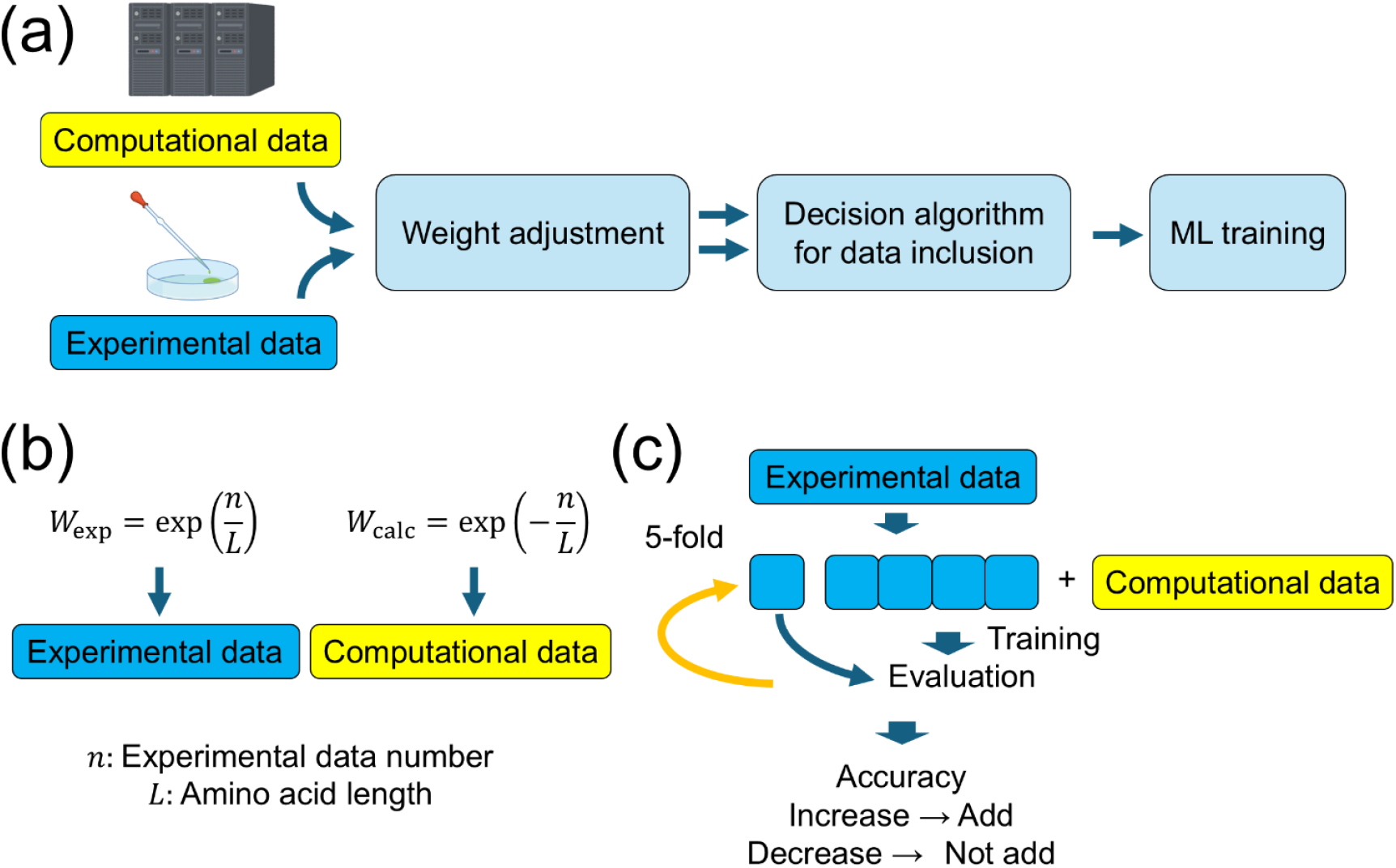
(a) Overview of the data-efficient protein mutational effect prediction method. Mutational effects are estimated by molecular simulation and pLM’s zero-shot prediction. These computational estimates are used as weak training data integrated with experimental training data for ML models. (b) The weight of weak training data relative to experimental training data are determined by the weight adjustment algorithm. (c) The decision algorithm determines whether or not to include weak training data based on cross validation.

Computational estimates of mutational effects involve errors in estimation as well as differences in scales and biases compared to experimental data. Thus, directly integrating them may cause a decrease in the prediction accuracy of resultant ML models. To prevent this, we developed the weight adjustment algorithm (Figure 1b) and the decision algorithm for data inclusion (Figure 1c). In the weight adjustment algorithm, the weight of computational estimates is reduced when the amount of experimental training data is large relative to the sequence length (Methods). In the decision algorithm, a subset of available experimental training data is used as validation data, and computational estimates are included only if their addition improves prediction accuracy (Methods).

To obtain computational estimates by Rosetta, we used different protocols depending on protein properties of interest (Methods). For protein abundance, the change in folding free energy upon mutations was calculated and converted to the protein abundance estimate using the biophysics-based equations. For binding affinity, the change in binding free energy with the ligand upon mutations was calculated and, together with the folding free energy, similarly converted to the binding affinity estimate. For enzymatic activity, due to the difficulty in simulating enzymatic reactions by Rosetta, we employed the folding free energy change upon mutations as used for protein abundance. To obtain computational estimates by ESM-2, we consistently used the log-likelihood ratio of mutant and wild-type sequences (Methods). The computational estimates by Rosetta and ESM-2 were further combined to calculate the hybrid score (Methods) that was used as weak training data in our method.

Our ML model takes as input a mutant sequence, calculates its feature values using ESM-2 (here used for embedding rather than zero-shot prediction), and feeds them into a downstream regression model to predict the mutational effect. The regression model was selected from support vector machine, random forest, gaussian process, and linear model for each dataset. We also tested models using amino acid descriptors for embedding instead of ESM-2-based embedding.

We evaluated our method on the datasets of experimentally measured mutational effects, which consist of various proteins and their properties including protein abundance, binding affinity, enzymatic activity, cytotoxicity, and fluorescence (Table 1). The datasets contain both single-residue and double-residue mutants. For each dataset, we first selected the regression model and its hyperparameters by nested cross validation on the whole experimental data (Supplementary Table 1).

**Table 1.**
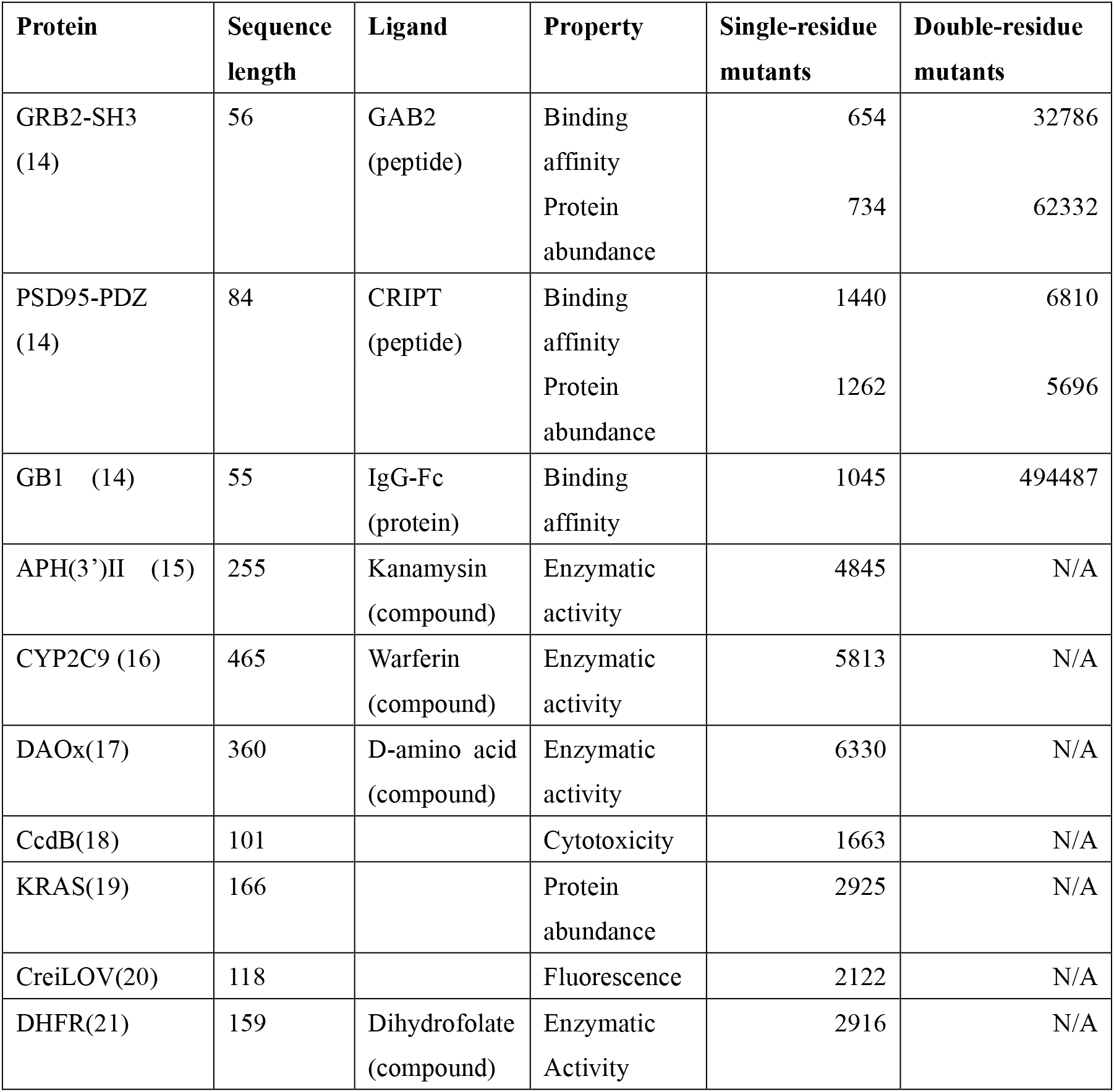
Benchmark dataset.

| Protein | Sequence length | Ligand | Property | Single-residue mutants | Double-residue mutants |
| --- | --- | --- | --- | --- | --- |
| GRB2-SH3<br>(14) | 56 | GAB2<br>(peptide) | Binding affinity | 654 | 32786 |
|  |  |  | Protein abundance | 734 | 62332 |
| PSD95-PDZ<br>(14) | 84 | CRIPT<br>(peptide) | Binding affinity | 1440 | 6810 |
|  |  |  | Protein abundance | 1262 | 5696 |
| GB1 (14) | 55 | IgG-Fc<br>(protein) | Binding affinity | 1045 | 494487 |
| APH(3')II (15) | 255 | Kanamycin<br>(compound) | Enzymatic activity | 4845 | N/A |
| CYP2C9 (16) | 465 | Warferin<br>(compound) | Enzymatic activity | 5813 | N/A |
| DAOx(17) | 360 | D-amino acid<br>(compound) | Enzymatic activity | 6330 | N/A |
| CcdB(18) | 101 |  | Cytotoxicity | 1663 | N/A |
| KRAS(19) | 166 |  | Protein abundance | 2925 | N/A |
| CreiLOV(20) | 118 |  | Fluorescence | 2122 | N/A |
| DHFR(21) | 159 | Dihydrofolate<br>(compound) | Enzymatic Activity | 2916 | N/A |

Then, the prediction accuracy on test data was compared before and after adding weak training data under the same model configuration to evaluate the impact of weak training data. We carefully devised the evaluation procedure so that the training process does not look at test data (Methods; Supplementary Figure 7). As the baseline, we also evaluated the prediction accuracy of Rosetta or ESM-2’s zero-shot prediction alone.

We also considered an end-to-end model using the ESM-2 encoder and a downstream regression network with all parameters trainable. However, this model resulted in low accuracy in data-scarce conditions (Supplementary Table 2), thus was not employed as our final model.

### Weak supervision improves prediction accuracy in data-scarce conditions

We first evaluated the impact of weak supervision using the single-residue mutant dataset for each protein (Figure 2). In most protein properties tested, incorporating weak training data improved the prediction accuracy compared to using experimental training data alone (Figure 2a-l). The improvement was more pronounced when the amount of experimental training data was small (e.g. <200). With the increase of experimental training data, the augmentation provided little additional benefit. When very few experimental training data were available (e.g. <50), predictions based solely on computational estimates (Rosetta or ESM-2) outperformed ML models trained with weak supervision. However, as the amount of experimental data increased, our method outperformed these computational estimates alone.

**Figure 2.**
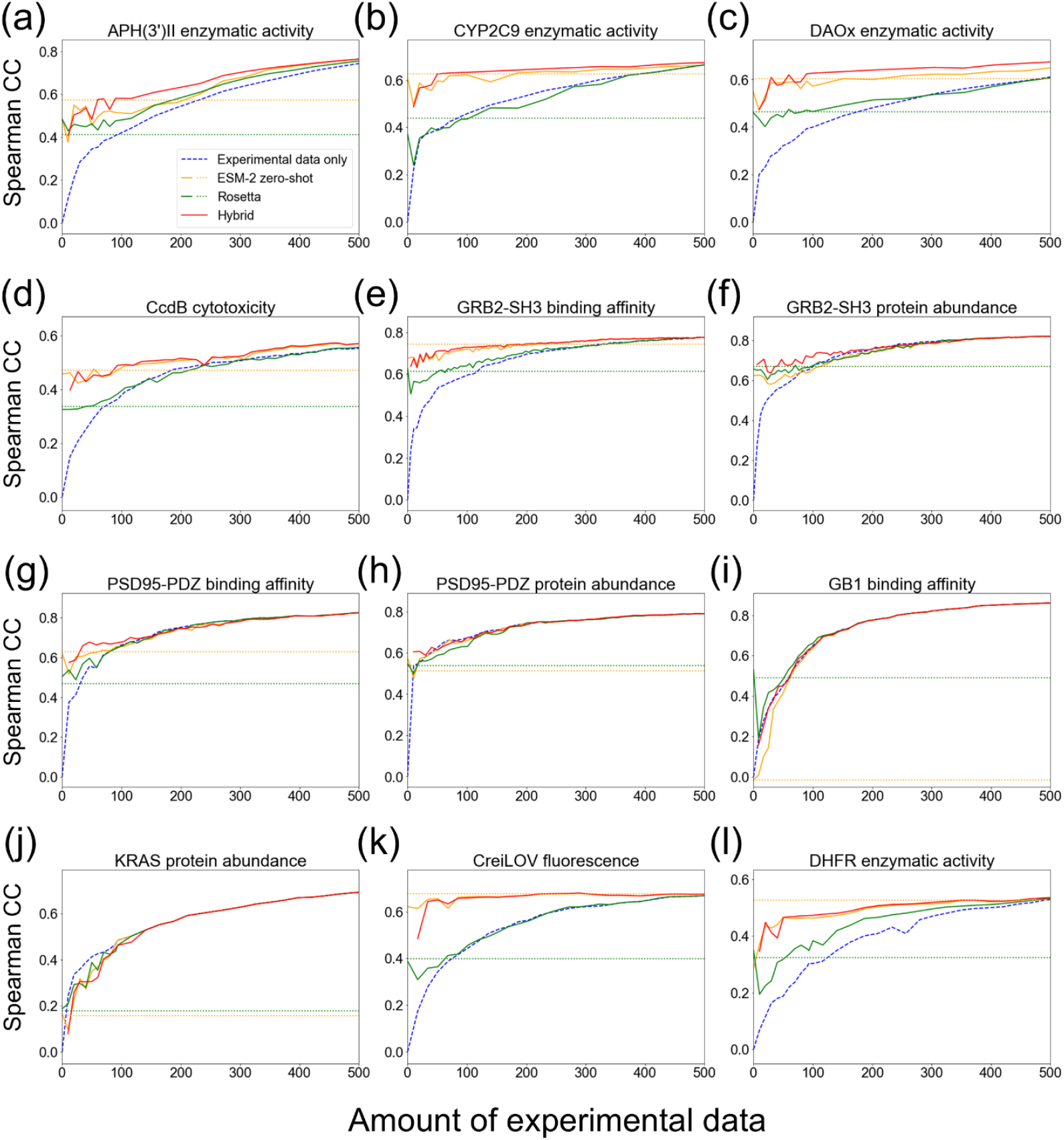
Impact of weak supervision with differing amounts of available experimental training data. Prediction accuracy measured by Spearman’s correlation coefficient (CC) was compared between ML models trained using experimental data only (dashed line), those integrated with computational estimates by Rosetta, ESM-2, and their hybrid score (solid lines). As the baseline, prediction accuracy obtained by computational estimates alone was shown (dotted lines). esm2_t6_8M_UR50D was used for embedding. Single-residue mutants were used for both training and test splits. (a) APH(3’)II enzymatic activity. (b) CYP2C9 enzymatic activity. (c) DAOx enzymatic activity. (d) CcdB cytotoxicity. (e) GRB2-SH3 binding affinity. (f) GRB2-SH3 protein abundance. (g) PSD95-PDZ binding affinity. (h) PSD95-PDZ protein abundance. (i) GB1 binding affinity. (j) KRAS protein abundance. (k) CreiLOV fluorescence. (l) DHFR enzymatic activity.

We also evaluated ML models using amino acid descriptors for embedding instead of ESM-2-based embedding (Supplementary Figure 1). As in the case of ESM-2-based embedding, the incorporation of weak training data improved prediction accuracy compared to the ML models trained solely with experimental training data. Our method also exceeded the computational estimates alone with the increase of experimental training data, with few exceptions of CYP2C9 enzymatic activity, GRB2-SH3 binding affinity, CreiLOV fluorescence, and DHFR enzymatic activity.

These results demonstrate that our method with weak supervision enhances prediction accuracy in data-scarce conditions while adapting to the different availability of experimental training data.

### Combining molecular simulation and pLM enables applicability to diverse protein properties

Molecular simulation based on biophysics and pLM pretrained on natural protein sequences may provide complementary estimates of mutational effects due to their different principles. To investigate this, we assessed prediction accuracy when using either Rosetta or ESM-2 alone as weak training data versus when combining both as the hybrid score (“Hybrid” in Figure 2). For GB1 binding affinity (Figure 2i), accuracy improved by adding Rosetta value, but not by ESM-2. For the other datasets, the hybrid score achieved the highest accuracy or at least comparable accuracy to Rosetta or ESM-2 alone. (Figure 2a-h, j-l).

When using amino acid descriptors for embedding, adding Rosetta value increased accuracy for GB1 binding affinity (Supplementary Figure 1i). For the other datasets, the hybrid score improved prediction accuracy the most or was comparable to Rosetta to ESM-2 alone (Supplementary Figure 1a-h, j-l).

Overall, these results indicate that combining molecular simulation with pLM-based zero-shot prediction enhances the applicability of our method across diverse protein properties.

### The weight adjustment algorithm and the decision algorithm for data inclusion provide robust prediction accuracy

To assess the impact of the weight adjustment algorithm and the decision algorithm for data inclusion, we conducted a comparison between using and not using these algorithms. We tested the two representative datasets, APH(3’)II enzymatic activity and PSD95-PDZ binding affinity (Figure 3). The weight adjustment algorithm substantially improved accuracy for APH(3’)II enzymatic activity (Figure 3a). However, accuracy increase was not notable with the decision algorithm (Figure 3a). To the contrary, the decision algorithm increased accuracy for PSD95-PDZ binding affinity, but weight adjustment did not (Figure 3b). By combining the two methods, accuracy was successfully increased in both datasets (Figure 3a, b). These results suggest that these algorithms are important for robust prediction accuracy, preventing potential negative impacts of weak training data.

**Figure 3.**
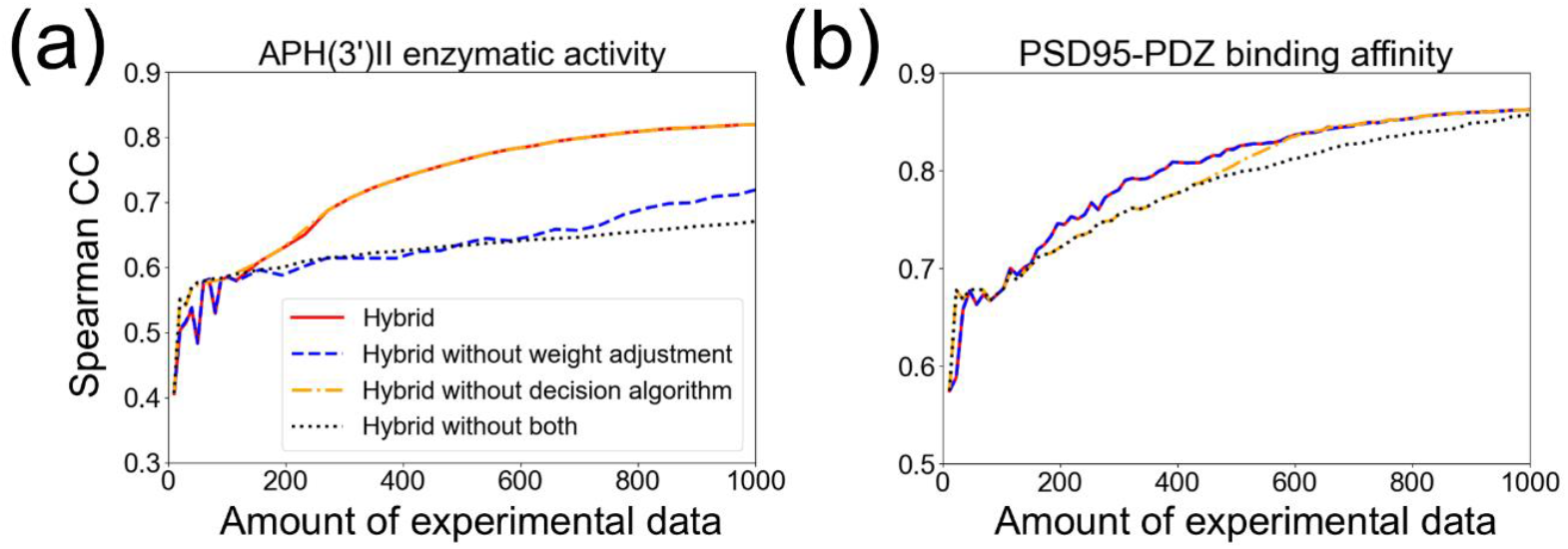
Ablation experiments for the weight adjustment algorithm and the decision algorithm for data inclusion. The prediction accuracy of ML model was compared among the hybrid score with both methods (Hybrid), without the weight adjustment algorithm, without the decision algorithm, and without both methods. esm2_t6_8M_UR50D was used for embedding. Single-residue mutants were used for both training and test splits. (a) APH(3’)II enzymatic activity. (b) PSD95-PDZ binding affinity.

### Predicting double-residue mutational effects using single-residue mutants as training data

In protein engineering, predicting the combinatorial effect of multiple-residue mutations is important to design functional proteins distant from the wild-type. Thus, we evaluated our method using double-residue mutants as test data while using single-residue mutants as training data. Remarkably, weak training data improved prediction accuracy in most of the protein properties compared to using experimental data alone (Figure 4). For PSD95-PDZ binding affinity and protein abundance (Figure 4c, d), the impact of weak supervision was relatively small while it still enhanced accuracy when experimental data was fewer than 50. For GB1 binding affinity (Figure 4e), accuracy improved the most by adding Rosetta value. For the other datasets, the hybrid score achieved the highest accuracy or at least comparable accuracy to that obtained by Rosetta or ESM-2. (Figure 4a-d).

**Figure 4.**
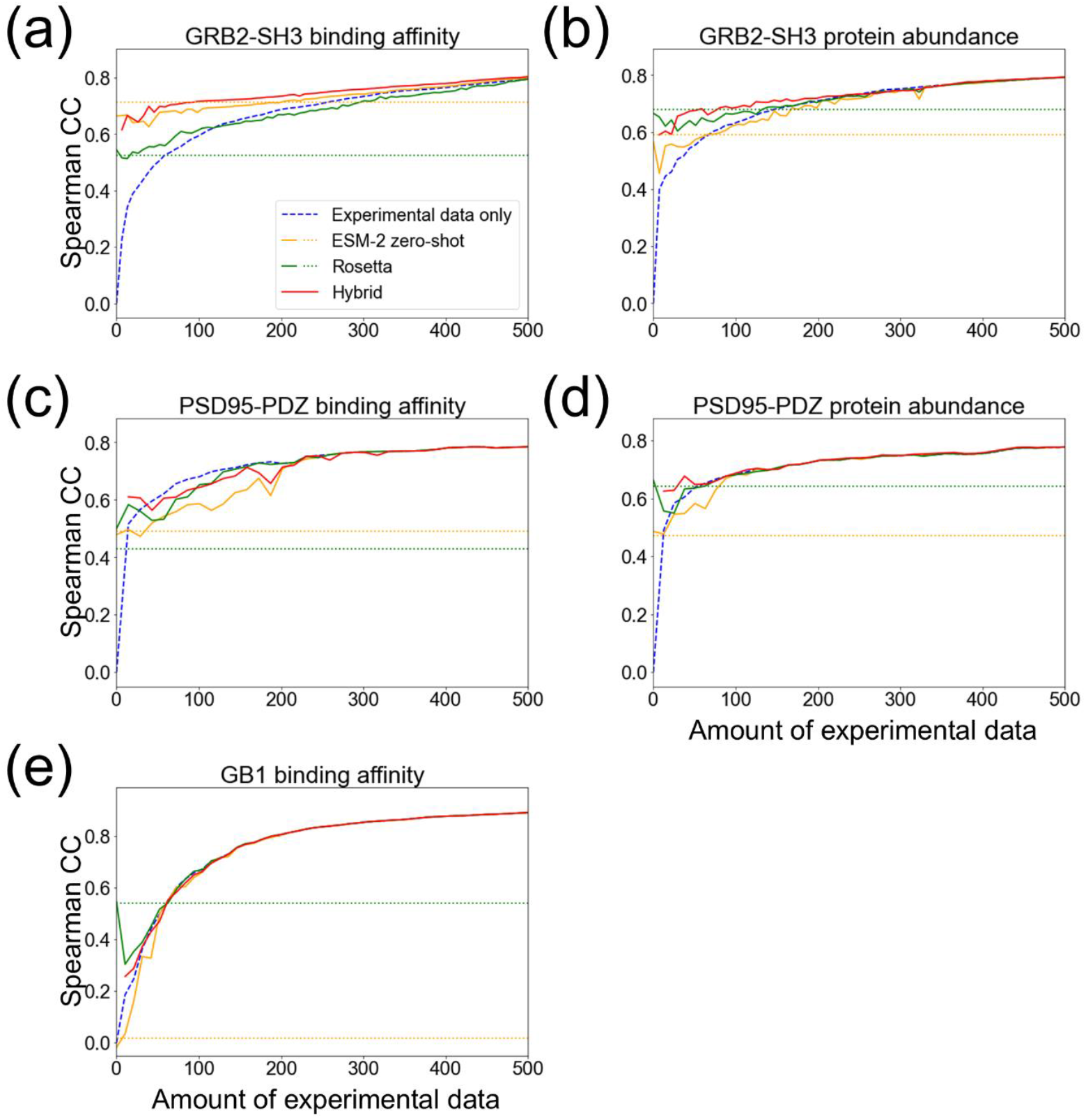
Prediction accuracy for double-residue mutational effects only using single-residue mutants as training data. ML models trained using experimental data only (dashed line), those integrated with computational estimates by Rosetta, ESM-2, and their hybrid score (solid lines) were compared. As the baseline, prediction accuracy obtained by computational estimates alone was shown (dotted lines). esm2_t6_8M_UR50D was used for amino acid embedding. (a) GRB2-SH3 binding affinity (b) GRB2-SH3 protein abundance. (c) PSD95-PDZ binding affinity. (d) PSD95-PDZ protein abundance. (e) GB1 binding affinity.

We also evaluated the models using amino acid descriptors for embedding (Supplementary Figure 2). As with ESM-2-based embedding, incorporating weak training data improved prediction accuracy compared to using experimental training data alone (Supplementary Figure 2a-e). For the datasets other than GRB2-SH3 binding affinity and GB1, the hybrid score improved accuracy the most (Supplementary Figure 2a-d). For GRB2-SH3 binding affinity (Supplementary Figure 2a), accuracy increased the most when adding ESM-2 value. For GB1, accuracy increased the most by adding Rosetta value (Supplementary Figure 2e).

The capability of our method to predict double-residue mutational effects could be less surprising if their corresponding single-residue mutations were available as experimental training data. For investigating to what extent this was the case, we calculated the fraction of double-residue mutants whose two mutations were both included as single-residue mutants in experimental training data. A double-residue mutant was regarded as “overlapped” if single-reside mutants were found at the same positions but the type of amino acids might differ (position level overlap), or the positions and the type of amino acids were the same (amino acid level overlap). We found that the position level overlap was relatively high (50-100%) while the amino acid level overlap was less than 10% at the experimental data size of 100 and 200. Thus, our method successfully predicted the effect of double-residue mutants even when their corresponding single-residue mutants were not available as experimental training data.

Overall, these results demonstrate that our method enhances prediction accuracy not only for single-residue mutational effects, but also double-residue mutations in data-scarce conditions.

### Weak supervision helps to find “hit” mutants

Protein engineering often aims to find “hit” mutants that have higher functions than the wild type. To evaluate our method in this scenario, we used another evaluation metric called hit rate (Supplementary Figure 3). The hit rate was defined as the fraction of hit mutants among the top 30 predicted mutants; we note that a similar metric was also used in a previous study (22). While the improvements by weak training data was less clear compared to the evaluations using Spearman’s correlation coefficient, we still observed the increased hit rate in several datasets (Supplementary Figure 3abdefj). For DFHR dataset, our method showed Spearman’s correlation coefficient lower than ESM-2 alone, but achieved higher performance in terms of the hit rate (Supplementary Figure 3k). These results indicate that weak supervision is a useful approach not only for improving prediction accuracy, but also for finding hit mutants in protein engineering.

### Comparison with the state-of-the-art mutational effect predictors

In addition to Rosetta and ESM-2’s zero-shot prediction, we further compared our method with the recently mutational effect prediction methods: SPIRED(23), GeoDDG(24), and DDGemb(25). Our method outperformed GeoDDG and DDGemb in most datasets (Supplementary Figure 4). On the other hand, we found that SPIRED uses some of the proteins in our datasets as its training data, thus the fair comparison with our method was difficult. Indeed, for these proteins (APH(3’)II, CYP2C9, PSD95-PDZ, CcdB, and DHFR), SPIRED tended to show higher or comparable accuracy to our method. Nevertheless, for the other proteins, our method consistently achieved better performance than SPIRED (Supplementary Figure 4cefijk). These results suggest that our method has the capability to perform better than these state-of-the-art methods, especially for external datasets not included in their training data.

## Discussion

In this study, we demonstrated that weak supervision using computational estimates by molecular simulation and pLMs can improve the accuracy of ML models for protein mutational effect prediction. We developed the weight adjustment algorithm and the decision algorithm for data inclusion, which successfully mitigated potential negative impacts caused by weak training data. Our results showed that integrating weak training data was particularly beneficial in data-scarce conditions where experimental data alone were insufficient to train accurate ML models. By combining Rosetta and ESM-2, our hybrid approach enhanced accuracy across diverse protein properties including protein abundance, binding affinity, and enzymatic activity.

Despite its effectiveness, weak supervision sometimes did not improve accuracy. This happened when computational estimates were quite inaccurate, even worse than ML models trained with small experimental data (e.g. Figure 2ij, Figure 4e). We speculate possible reasons for inaccurate computational estimates by Rosetta and ESM-2 as follows. Rosetta-based estimates possibly become inaccurate for proteins with e.g. complex conformational changes that are not well-represented in molecular simulation. In addition, EMS-2’s zero-shot prediction may not correlate with mutational effects for proteins with limited representation in its pre-training dataset.

Rosetta and ESM-2 do not directly compute protein properties, especially those such as cytotoxicity. Nonetheless, these computational estimates showed a correlation with diverse protein properties, thereby serving as weak training data in our method. We attribute this fact to that Rosetta and ESM-2 may capture a protein’s “general fitness”. Rosetta computes the folding free energy; since structure folding is a prerequisite for a protein to exhibit its function, it may correlate with diverse protein properties measured by experiments. Similarly, ESM-2 computes the likelihood (protein-likeness) learned from natural proteins; thus, it may correlate with diverse protein properties observed in nature. For example, in our cytotoxicity dataset, since CcdB’s function in nature is a bacterial toxin, its folding free energy (Rosetta) and the likelihood (ESM-2) may correlate with its cytotoxicity. Based on this interpretation, we speculate that Rosetta and ESM-2 may not provide good weak training data when the protein property of interest is different from its natural function, which sometimes occur in protein engineering e.g. trying to enhance enzymatic activity for unnatural substrates etc. While this point was not systematically investigated in the present study, it will be a possible future direction.

Compared to previous studies, our approach extends the applicability of computationally augmented mutational effect prediction. A notable prior work, mGPfusion, leveraged Rosetta-based estimates for weak supervision but was limited to thermostability prediction. Our method generalizes this concept by incorporating pLM-based estimates, allowing for the prediction of a wider range of protein properties, including binding affinity and enzymatic activity. In addition, we newly introduced the weight adjustment algorithm and decision algorithm for data inclusion, preventing unreliable computational estimates from negatively impacting model performance.

It is important to give a guideline about the expected amount of experimental training data for using our method. Looking at data-scarce conditions with less than 100 experimental training data (Supplementary Figures 5 and 6), the improvement by weak supervision was observed in many datasets before the experimental data size reached 100. Therefore, we propose to use 50-100 experimental data as an initial configuration for our method.

Overall, this study highlights the potential of weak supervision to address data scarcity in mutational effect prediction. By combining molecular simulation and pLM’s zero-shot prediction, we provide a widely applicable framework for improving ML-based protein engineering and pathogenicity prediction. Future work could explore additional computational estimation methods such as other AI-based methods (26, 27) or biophysics-based methods(28, 29). The ability to improve prediction accuracy without increasing experimental costs has significant benefits for accelerating protein design and understanding genotype-phenotype relationships.

## Methods

## Benchmark datasets

We collected benchmark datasets consisting of experimentally measured mutational effects on diverse protein properties, including protein abundance, binding affinity, enzymatic activity, cytotoxicity, and fluorescence (Table 1). These proteins originate from various species including human, other eukaryotes, and prokaryotes: GRB2-SH3, PSD95-PDZ, KRAS, and CYP2C9 from human; APH3 from *Klebsiella pneumoniae*; GB1 from *Streptococcus* sp. group G; DAOx from *Rhodotorula gracilis*; CcdB and DHFR from *Escherichia coli*; and CreiLOV from *Chlamydomonas reinhardtii*. The binding affinities of GRB2-SH3, PSD95-PDZ, and GB1, as well as the protein stabilities of GRB2-SH3, PSD95-PDZ, and KRAS, were measured using yeast growth. DHFR enzymatic activity and CcdB cytotoxicity were measured using *E. coli* growth. The enzymatic activities of APH(3’)II and CYP2C9 were assessed based on drug resistance in *E. coli* and yeast, respectively. DAOx enzymatic activity was assessed with fluorescent signal. CreiLOV fluorescence was measured with fluorescence-activated cell sorting. To systematically evaluate the impact of weak supervision under varying data availability, we selected datasets generated from high-throughput experiments, each containing at least 500 single-residue mutants per protein.

Mutational effect values were extracted from the original publications as follows. For the APH(3’)II enzymatic activity dataset, we used measurements obtained under the 1:8 dilution condition of the kanamycin ligand(15). The original study reported *ΔMut* and *ΔAA*, representing the changes in observed mutant frequencies at each site and for each amino acid, respectively. We defined the mutational effect for each amino acid substitution at each site as log(*ΔMut\**Δ*AA*). For the CYP2C9 enzymatic activity dataset, we directly used the site- and amino acid-specific mutational effects reported in the original study(16). For the GRB2-SH3, PSD95-PDZ, and GB1 datasets, mutational effects on protein abundance and binding affinity were obtained from a previous study(14). In addition, this study also reported folding free energy and binding free energy changes upon mutation, which were computed using biophysics-based conversion equations. For single-residue mutants, we included only those with both experimentally measured mutational effects and corresponding free energy changes available. All double-residue mutants were retained without filtering. For KRAS, the mutational effect and folding free energy dataset was obtained and constructed with the same way(19). For CcdB, DHFR, and CreiLOV, the mutational effect and folding free energy datasets were obtained from Proteingym(30) (CcdB: CCDB_ECOLI_Tripathi_2016, DHFR: DYR_ECOLI_Nguyen_2023, and CreiLOV: PHOT_CHLRE_Chen_2023).

## Molecular simulation

We used Rosetta ddG monomer (31) to calculate the change in folding free energy upon mutations (*ΔΔG*_**f**_). Rosetta Flex ddG (32) was used to calculate the change in binding free energy with the ligand upon mutations (*ΔΔG*_**b**_). We used the tertiary structure of each protein from PDB (33) (PDB IDs, GRB2-SH3: 2VWF(34), PSD95-PDZ: 1BE9(35), GB1: 1FCC(36), APH(3’)II: 1ND4(37), and CYP2C9: 1R9O(38), KRAS: 6VJJ(39), CcdB: 3VUB(40), DAOx: 1C0P(41), and DHFR: 1RX2(42)).

We used the same PDB entry as in the literature reporting each mutational effect dataset. For CreiLOV protein, we used a structure predicted by AlphaFold2(43) since no experimental structure was available. PyMOL(44) was used for correcting mutated residue in experimental structures. Similarly, Modeller(45) was used to predict structure of missed residue in the middle of the chain. For *ΔΔG*_**f**_ calculations, ligands were removed from the structure. In the ddG monomer calculations, high-resolution backrub-based protocol 16, which allows backbone conformational freedom and optimize the rotamers at all residues, was used for GRB2-SH3, PSD95-PDZ, GB1, and APH(3’)II, while the low-resolution side-chain flexibility based protocol 3, which only allows sidechain flexibility and optimizes only the rotamers for the residues in the neighborhood of the mutation, was applied to CYP2C9, KRAS, CcdB, DAOx, DHFR, and CreiLOV.

To obtain the computational estimates of mutational effects from Rosetta-derived *ΔΔG*_**f**_ and *ΔΔG*_**b**_, we applied the biophysics-based conversion equations inspired by(14):

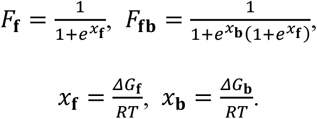

*ΔG*_**f**_ and *ΔG*_**b**_ represent Gibbs free energies of folding and binding, respectively, and *R* is gas constant and *T* is the temperature in Kelvin. As proposed in(14), *F*_**f**_ indicates the probability that the protein is in the folded state while *F*_**fb**_ indicates the probability that the protein is both in the folded state and bound to the ligand. Therefore, we used *F*_**f**_ and *F*_**fb**_ as the computational estimates for protein abundance and binding affinity, respectively. For enzymatic activity, cytotoxicity, and fluorescence, we used *F*_**f**_ as the computational estimates. This choice was motivated by the difficulty of simulating these protein properties with molecular simulation and the observation that folding stability was often strongly correlated with these protein properties.

## Zero-shot prediction with pLM

We used ESM-2 to perform zero-shot prediction and obtain computational estimates of mutational effects(12, 13). Specifically, we employed the 3-billion parameter model (esm2_t36_3B_UR50D) and adopted the wild-type marginal probability scheme for scoring mutant sequences relative to the wild type.

## Hybrid score

In addition to using Rosetta- and ESM-2-derived estimates independently, we also constructed a hybrid score by linearly combining the two values. This approach aims to improve robustness by integrating predictions derived from different principles: Rosetta being biophysics- and structure-based, and ESM-2 being deep learning- and sequence-based. The hybrid score was defined as follows:

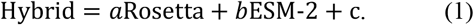

The coefficients *a, b*, and *c* were optimized using linear regression between the hybrid score and experimental mutational effects in training data (Supplementary Figure 7). This hybrid score was then used as weak training data in the same manner as the individual estimates.

## Scaling

Since the calculated values of *ΔΔG*_**f**_, *ΔΔG*_**b**_, *F*_**f**_, *F*_**fb**_, and ESM-2 values may differ in scales from their corresponding experimental values, we applied linear scaling using mutants in training data. For the GRB2-SH3, PSD95-PDZ, and GB1 datasets, *ΔΔG*_**f**_ was scaled using the following equation:

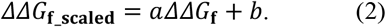

The coefficients *a* and *b* were determined by fitting *ΔΔG*_f_**scaled**_ to folding free energy available in training data (Supplementary Figure 7). After scaling, the change in folding free energy (*ΔΔG*_**f**_) was transformed into folding free energy (*ΔG*_**f**_), which was used for the calculation of *F*_**f**_ and *F*_**fb**_. The same scaling procedure was applied to *ΔΔG*_**b**_, *F*_**f**_, and *F*_**fb**_, and ESM-2 values. For the APH(3’)II and CYP2C9 datasets, where folding free energy and binding free energy changes were not reported in the original studies, we performed empirical scaling. Specifically, *ΔΔG*_**f**_ and *ΔΔG*_**b**_ were normalized to have a mean of 0 and a standard deviation of 3.0.

## Weight adjustment algorithm

To mitigate the potential negative impact of inaccuracies in computational estimates, we applied different weights to experimental training data and weak training data during model training. The weights were defined as follows:

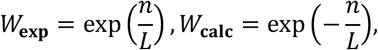

where *n* is the number of experimental training data, and *L* is the number of residues in the protein sequence where mutations can occur. This formulation increases the relative importance of experimental data as more experimental measurements become available while reducing the influence of computational estimates.

The rationale behind this approach is illustrated in Supplementary Figure 8. When experimental data are scarce, many regions in the sequence space are only covered by computational estimates (yellow area). However, as experimental data accumulate, the regions influenced by high-quality data (dashed circle) expand. Without proper reweighting, computational estimates may interfere with learning in these regions, leading to reduced accuracy. The weight adjustment algorithm addresses this by shifting learning emphasis away from computational estimates and toward experimental data in these regions, while still leveraging computational estimates in unexplored areas. This strategy balances model performance across the sequence space and contributes to overall prediction accuracy.

## Decision algorithm for data inclusion

In addition to the weight adjustment algorithm, we developed a decision algorithm to determine whether computational estimates should be included in model training.

The available experimental data were split into five subsets for cross validation. In each fold, four subsets were used for training, and the remaining subset was used as validation data. Computational estimates were added to the training set, and a model was trained on the combined dataset. The prediction accuracy on the validation set was then compared to that obtained using only experimental data. If the inclusion of computational estimates reduced validation accuracy, the algorithm rejected the addition of computational estimates as weak training data (Figure 1c).

In cases where experimental data was too small and the validation accuracy resulted in NaN, the algorithm defaulted to including computational estimates. This ensures that weak supervision can still be applied in data-scarce settings, where experimental guidance is limited.

## Machine learning

For each mutant, an amino acid sequence was embedded using either ESM-2 embedding (esm2_t6_8M_UR50D) or predefined amino acid descriptors. Letting *L* denotes the sequence length, the matrix of shape (*L*, 320) was generated by esm2_t6_8M_UR50D, which was flattened along the *L*-axis into a 320*L*-dimensional vector. The descriptor was selected from among BLOSUM-based features(46), FASGAI(47), MS-WHIM(48), T-scale(49), ST-scale(50), Z-scale(51), VHSE(52), and ProtFP(53). Regression models were selected from random forest(54), support vector machine(55), linear regression, and Gaussian process regression(56). For each protein property, the embedding method, regression model, and its hyperparameters were first selected by nested cross-validation on the dataset of all single-residue mutants (Supplementary Table 1). These configurations were then fixed and consistently used for the benchmark tests to evaluate the impact of data augmentation.

All model training and prediction were performed using the scikit-learn library in Python(57). Due to large computational costs, random forest was excluded for the APH(3’)II, CYP2C9, DHFR, DAOx, KRAS, CreiLOV, and DHFR datasets when ESM-2 embeddings were used.

We also considered end-to-end neural networks as model candidates. Specifically, we constructed a model using the ESM-2 encoder and a downstream regression network with all parameters trainable. However, this model resulted in lower accuracy than the above models in data-scarce conditions (Supplementary Table 2), thus was not employed as our final model.

## Training and evaluation

For evaluations on single-residue mutants, we first randomly selected 20% of the mutants as hold-out test data. The remaining 80% served as the pool from which training data were sampled (see below). This random split was repeated 10 times with different random seeds, and the average performance was reported.

For evaluations on double-residue mutants, all single-residue mutants in each dataset were used as the training data pool. All available double-residue mutants were used as test data, except for the GB1 dataset with ESM-2 embeddings where 10,000 mutants were randomly selected for each evaluation due to large computational costs.

To simulate different levels of experimental data availability, we constructed training sets by randomly sampling *n*% (*n* = 1, 2, …, 99) from the above training data pool (Supplementary Figure 7). The remaining data were labeled with computational estimates from Rosetta and ESM-2, which were used for data augmentation. For the sampled training data, Rosetta and ESM-2 values were also computed and used to determine the coefficients for the hybrid score (Equation 1) and scaling function (Equation 2). Prediction accuracy was evaluated on the hold-out test data using Spearman’s correlation coefficient, with and without data augmentation. Importantly, test data were never used during the model training and the fitting of the hybrid score and scaling functions to prevent data leakage. This procedure was repeated 10 times per sampling rate *n* using different random seeds, and the average performance was reported.

For the APH(3’)II, CYP2C9, DHFR, DAOx, and KRAS datasets, which contain a large number of mutants, we also evaluated under training data points fewer than 100 to more closely examine performance in low-data conditions.

To assess the performance of Rosetta and ESM-2 computational estimates alone (i.e., without ML models), we used the same test data as in the ML evaluation for single-residue mutants. For double-residue mutants, 5,000 variants were randomly sampled as test data. From these, 500 variants were randomly selected for 10 times to calculate the average performance. No scaling (Equation 2) was applied in these baseline evaluations.

## Data Availability

The source code and the benchmark datasets used in this study are available at https://github.com/Teppei-Deguchi/prj-weak-supervision

## Supporting Information

Supplementary Figures 1-8 and Supplementary Table 1-3.

## Author Contributions

TD and YS developed the method and wrote the paper. TD wrote the code and performed the computational experiments. NSAB helped model implementation. YK, SI, and KK participated in the data interpretation. All authors read and approved the final version of the manuscript.

## Acknowledgements

This work was supported by JSPS KAKENHI (22H03691), AMED grant (JP19ak0101122), NEDO grant (JPNP14004), JACI Prize for Encouraging Young Researcher (to YS), and JST SPRING (JPMJSP2108) (to TD). The computations were partly performed on the NIG supercomputer at ROIS National Institute of Genetics, the ABCI supercomputer at AIST, and the SQUID supercomputer at Cybermedia Center, Osaka University through the HPCI System Research Project (hp230057, hp240075). TD gratefully acknowledges financial support from Hattori international scholarship foundation and G-7 scholarship foundation.

