## Supplementary Information for "Data-efficient protein mutational effect prediction with weak supervision by molecular simulation and protein language models"

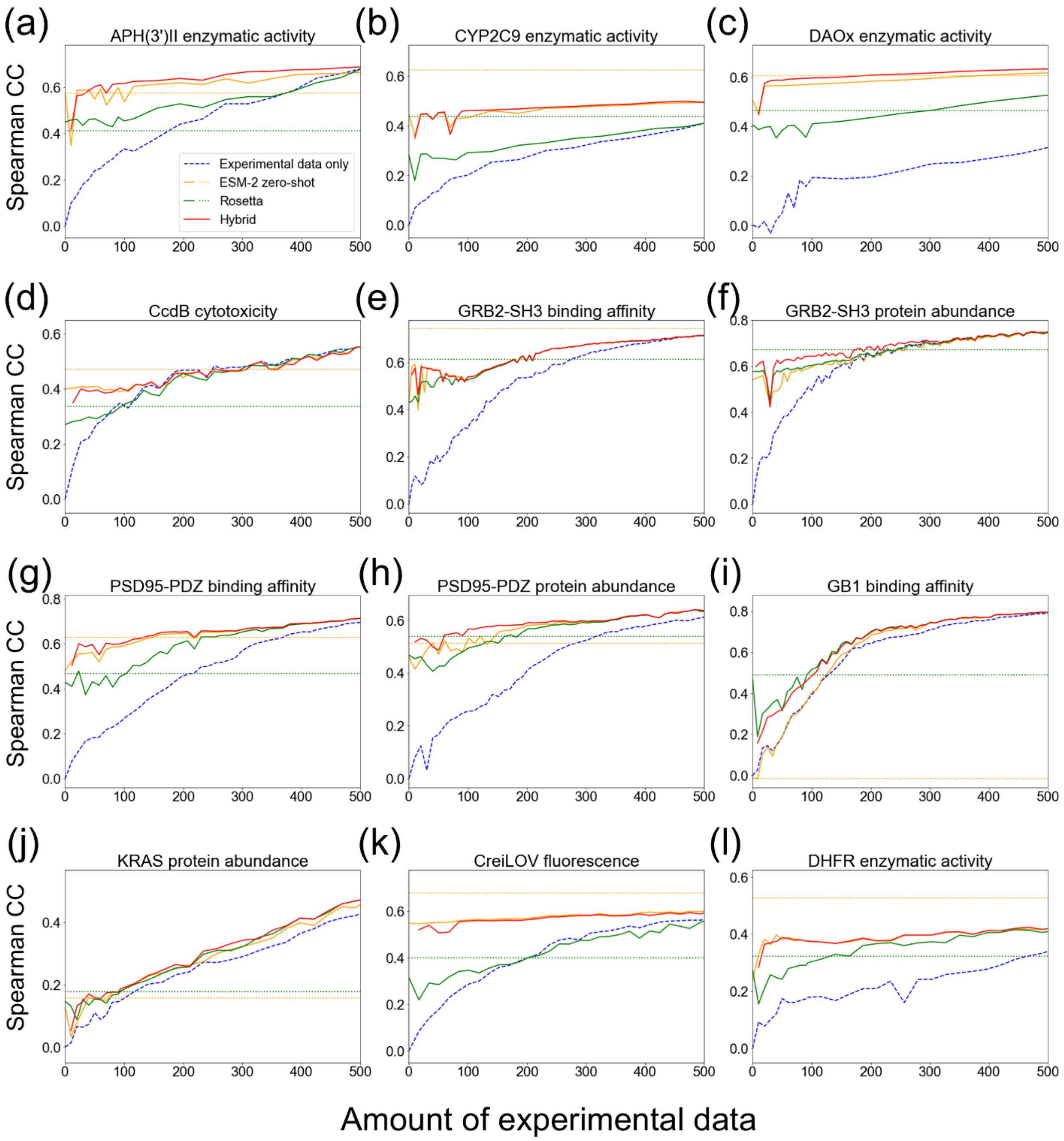


Supplementary Figure 1. Prediction accuracy measured by Spearman's correlation coefficient (CC) was compared between ML models trained using experimental data only (dashed line), those integrated with computational estimates by Rosetta, ESM-2, and their hybrid score (solid lines). As the baseline, prediction accuracy obtained by computational estimates alone was shown (dotted lines). Amino acid descriptors were used for embedding as shown in Supplementary Table 1. (a) APH(3')II enzymatic activity. (b) CYP2C9 enzymatic activity. (c) DAOx enzymatic activity. (d) CcdB cytotoxicity. (e) GRB2-SH3 binding affinity. (f) GRB2-SH3 protein abundance. (g) PSD95-PDZ binding affinity. (h) PSD95-PDZ protein abundance. (i) GB1 binding affinity. (j) KRAS protein abundance. (k) CreiLOV fluorescence. (l) DHFR enzymatic activity.

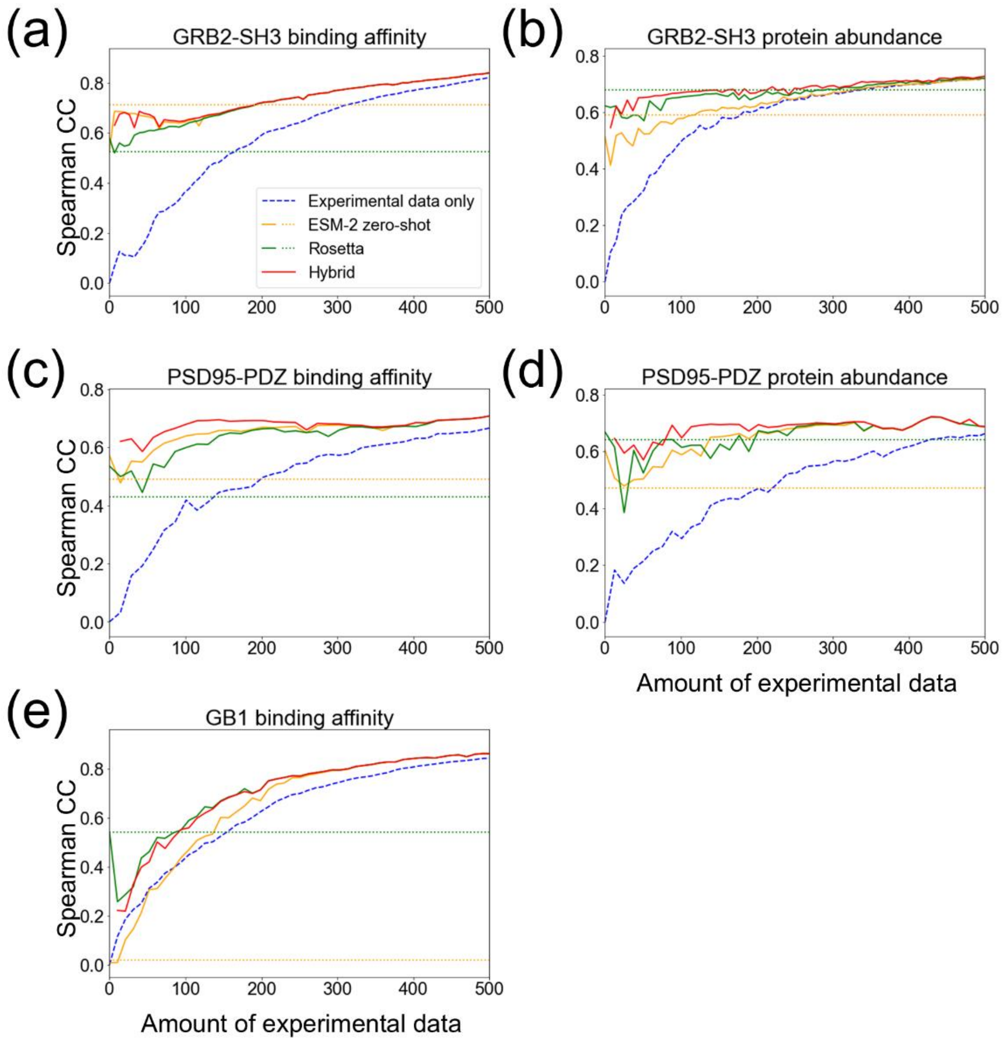


Supplementary Figure 2. Prediction accuracy for double-residue mutational effects only using single-residue mutants as training data. ML models trained using experimental data only (dashed line), those integrated with computational estimates by Rosetta, ESM-2, and their hybrid score (solid lines) were compared. As the baseline, prediction accuracy obtained by computational estimates alone was shown (dotted lines). Amino acid descriptors were used for embedding as shown in Supplementary Table 1. (a) GRB2-SH3 binding affinity (b) GRB2-SH3 protein abundance. (c) PSD95-PDZ binding affinity. (d) PSD95-PDZ protein abundance. (e) GB1 binding affinity.

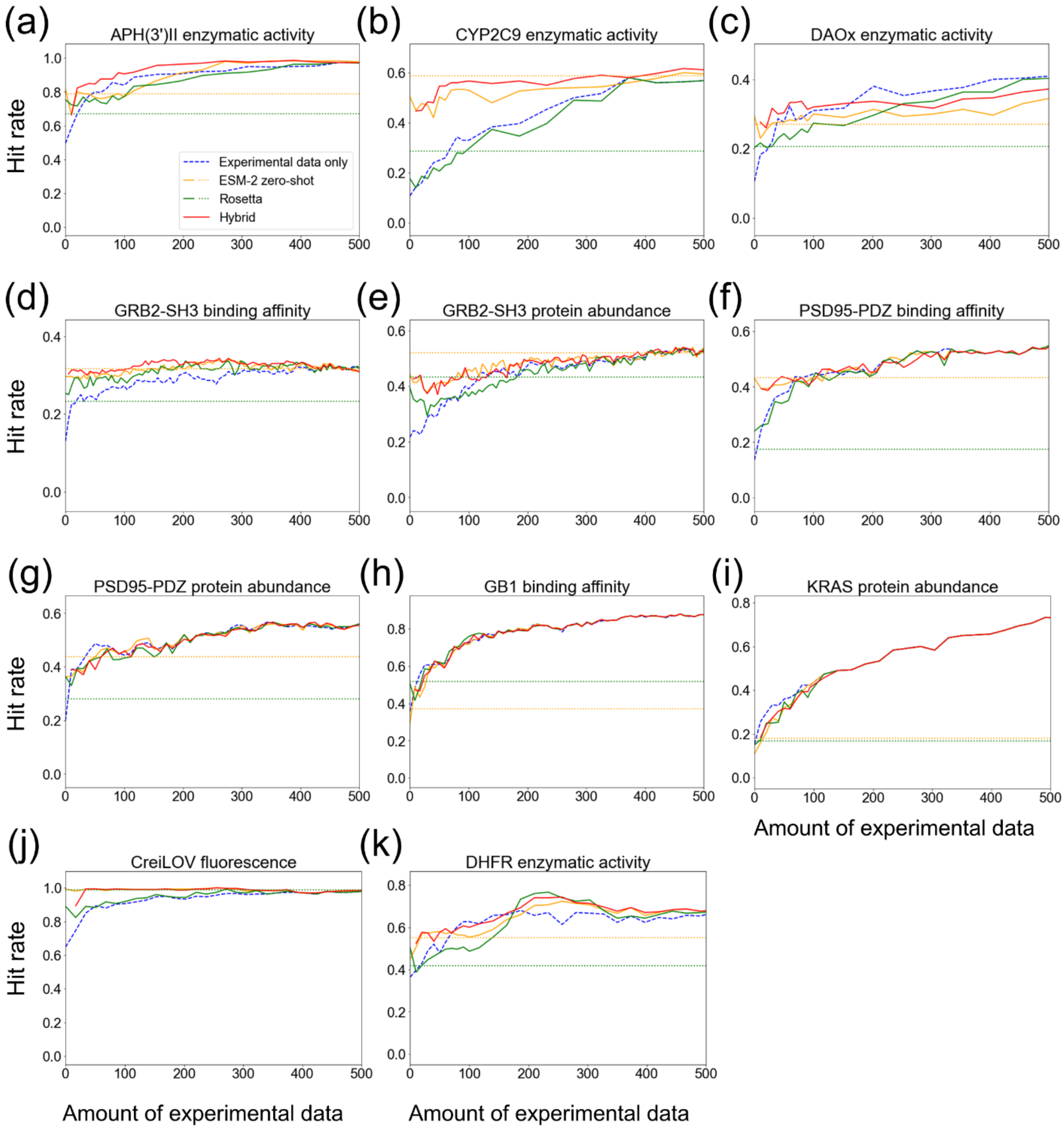


Supplementary Figure 3. Evaluation based on the hit rate. Hit mutants are defined as mutants that have higher functions than the wild type. The hit rate is the fraction of hit mutants among the top 30 predicted mutants. The average hit rate of 10-times experiments was shown. CcdB dataset was not used since the wild-type value was not provided in the dataset. For the other explanations, see the caption of Figure 2.

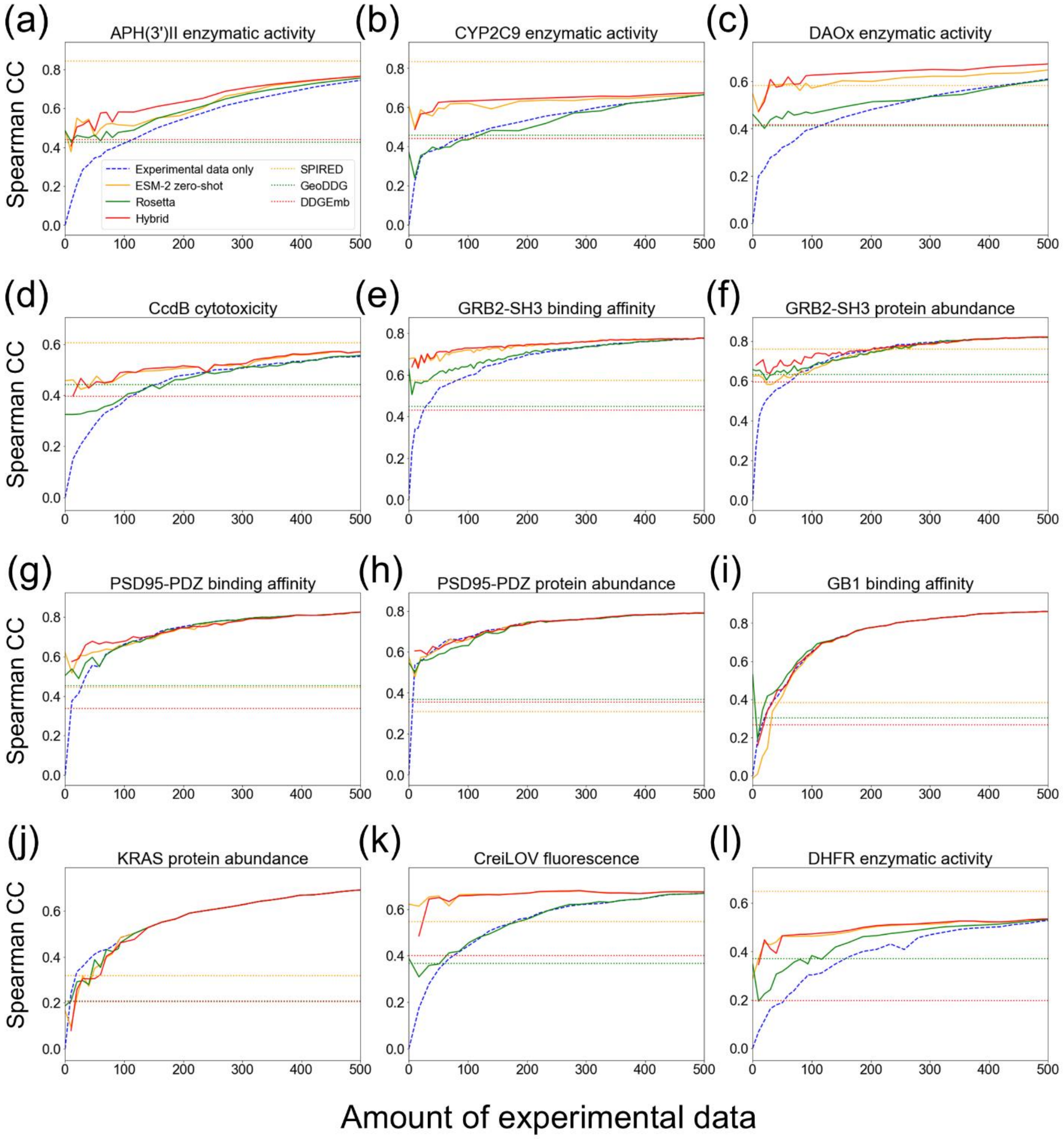


Supplementary Figure 4. Prediction accuracy measured by Spearman's correlation coefficient (CC) was compared to the state-of-the-art mutational effect predictors: SPIRED, GeoDDG, and DDGemb. esm2_t6_8M_UR50D were used for embedding as shown in Supplementary Table 1. Note that SPIRED uses the mutational effect data of APH(3')II, CYP2C9, PSD95-PDZ, CcdB, and DHFR as its training data. (a) APH(3')II enzymatic activity. (b) CYP2C9 enzymatic activity. (c) DAOx enzymatic activity. (d) CcdB cytotoxicity. (e) GRB2-SH3 binding affinity. (f) GRB2-SH3 protein abundance. (g) PSD95-PDZ binding affinity. (h) PSD95-PDZ protein abundance. (i) GB1 binding affinity. (j) KRAS protein abundance. (k) CreiLOV fluorescence. (l) DHFR enzymatic activity.

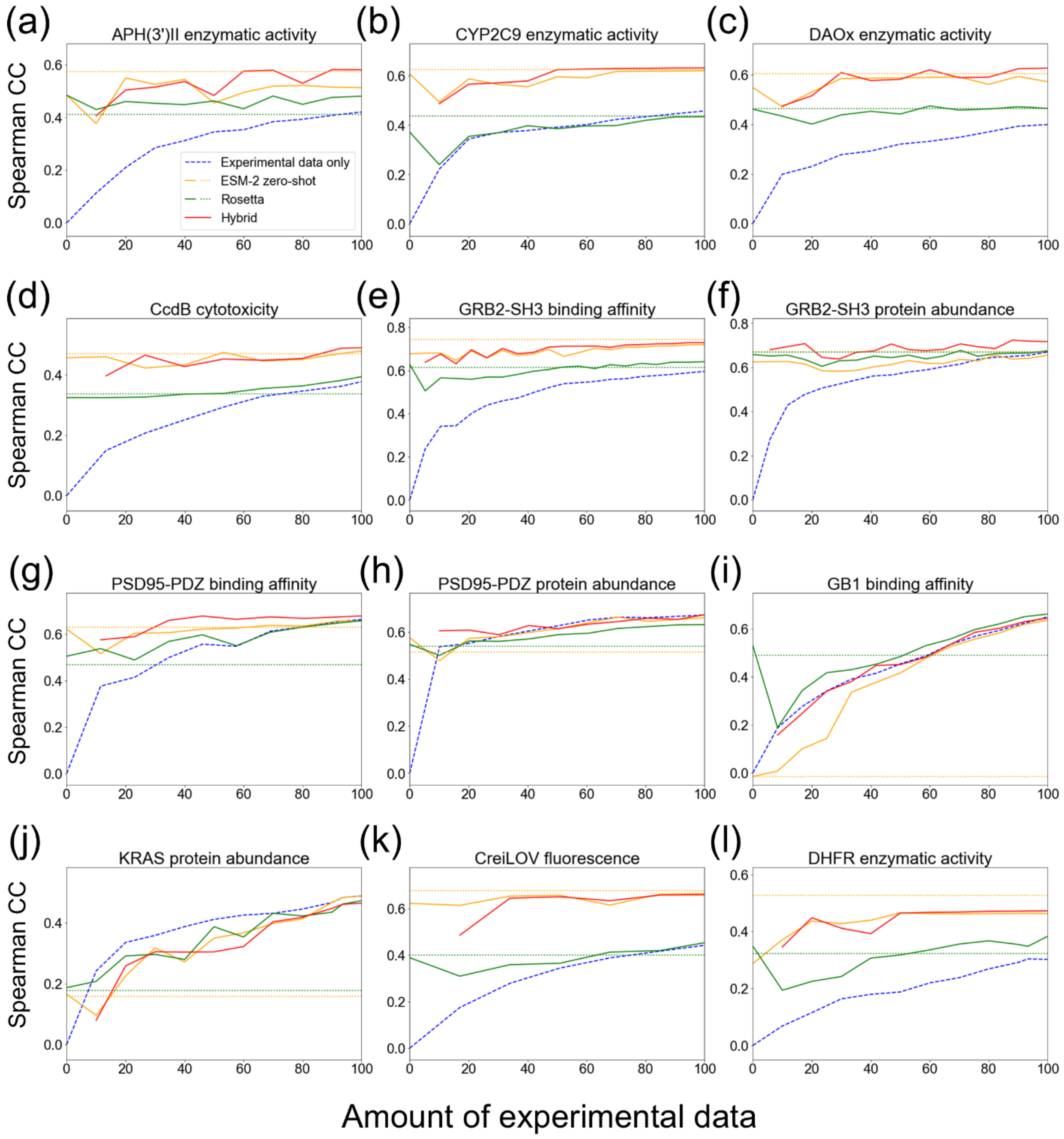


Supplementary Figure 5. Magnified version of Figure 2 focusing on small data regimes (<100). See the caption of Figure 2 for the other explanations.

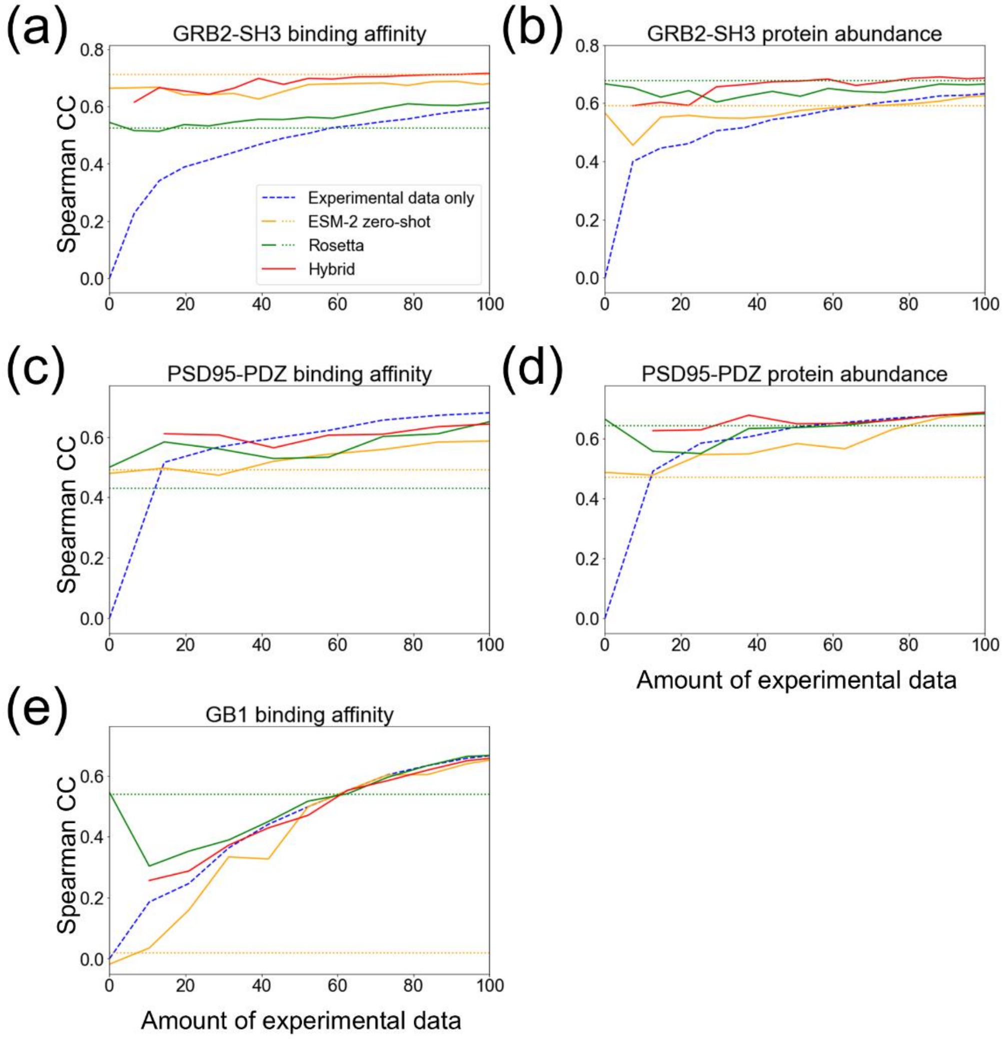


Supplementary Figure 6. Magnified version of Figure 4 focusing on small data regimes (<100). See the caption of Figure 4 for the other explanations.

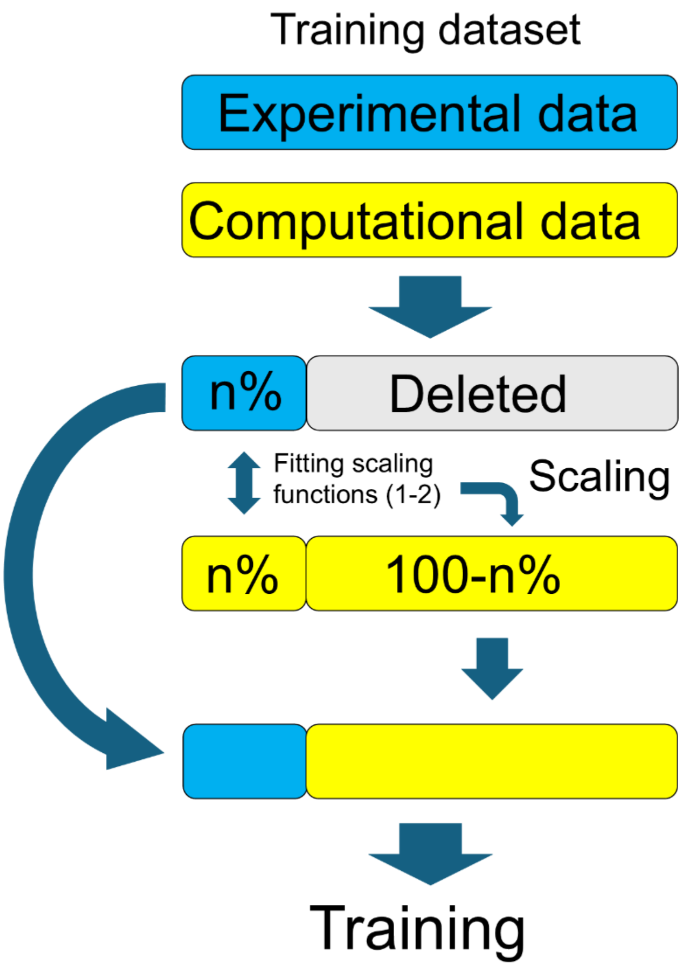


Supplementary Figure 7. The process of data augmentation. Mutant sequences in experimental training data are labeled with computational estimates by Rosetta and ESM-2, which are used for data augmentation and fitting scaling functions.

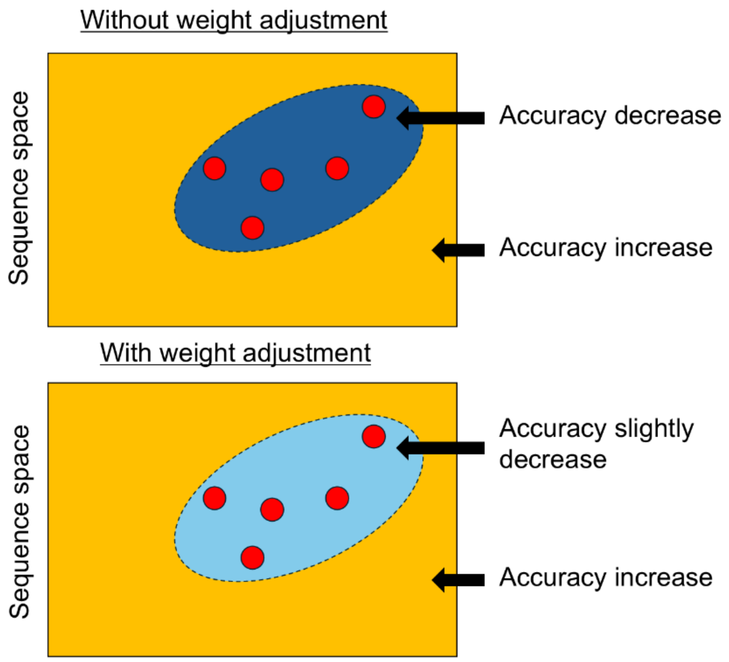


Supplementary Figure 8. Conceptual illustration of how weight adjustment works in sequence space to prevent accuracy degradation. We assume augmented data are broadly distributed in sequence space as shown in yellow, and red points represent experimental data points. The dashed circle represents the region where rich experimental training data are available.

Supplementary Table 1. ML configurations. Feature, ML model, and parameters were selected with nested cross validation. Support vector regression (SVR); random forest regression (RFR). All SVR parameters were selected from powers of 10. RFR parameter n_estimators is selected from multiples of 500, such as 500, 1000, and 1500. For CPY2C9 with ESM-2 embedding, C=10, ε= 1e-6 – 1e-2, γ= 1e-8 – 1e-5 was excluded due to the computational cost.

| **Protein** | **Functional value** | **Feature** | **ML model** | **Parameters** |
|---|---|---|---|---|
| GRB2-SH3 | Binding affinity | BLOSUM | SVR | C=0.1, ε=0.01, γ=0.001 |
| | | esm2_t6_8M_UR50D | SVR | C=1000, ε=0.0001, γ=0.0001 |
| | Abundance | FASGAI | RFR | n_estimators=500 |
| | | esm2_t6_8M_UR50D | SVR | C=1000, ε=1e-06, γ=1e-05 |
| PSD95-PDZ | Binding affinity | VHSE | SVR | C=1, ε=0.01, γ=0.001 |
| | | esm2_t6_8M_UR50D | SVR | C = 1000, ε = 1e-05, γ = 1e-05 |
| | Abundance | FASGAI | SVR | C=1, ε=0.01, γ=0.001 |
| | | esm2_t6_8M_UR50D | SVR | C=10, ε=0.01, γ=1e-05 |
| GB1 | Binding affinity | VHSE | SVR | C=10.0, ε=0.01, γ=0.001 |
| | | esm2_t6_8M_UR50D | SVR | C=10, ε=0.001, γ=1e-05 |
| APH(3')II | Activity | FASGAI | RFR | n_estimators=1000 |
| | | esm2_t6_8M_UR50D | SVR | C=1, ε=1e-06, γ=1e-06 |
| CPY2C9 | Activity | BLOSUM | RFR | n_estimators=1500 |
| | | esm2_t6_8M_UR50D | SVR | C=100, ε=1e-05, γ=1e-06 |
| DAOx | Activity | VHSE | SVR | C=1, ε=0.01, γ=1e-04 |
| | | esm2_t6_8M_UR50D | SVR | C=1, ε=0.001, γ=1e-05 |
| CcdB | Activity | Z-scale | RFR | n_estimators=500 |
| | | esm2_t6_8M_UR50D | SVR | C=100, ε=0.0001, γ=1e-05 |
| KRAS | Abundance | VHSE | SVR | C=1, ε=0.01, γ=0.001 |
| | | esm2_t6_8M_UR50D | SVR | C=10, ε=0.0001, γ=1e-06 |
| CreiLOV | Activity | VHSE | RFR | n_estimators=2500 |
| | | esm2_t6_8M_UR50D | SVR | C=10, ε=1e-06, γ=1e-06 |
| DHFR | Activity | ST-scale | SVR | C=0.1, ε=1e-06, γ=1e-04 |
| | | esm2_t6_8M_UR50D | SVR | C=0.1, ε=1e-07, γ=1e-05 |

Supplementary Table 2. Comparison between SVR and ESM-2 supervised fine-tuning (ESM-2 SFT). The amount of experimental training data was fixed as 100. esm2_t6_8M_UR50D was used as the encoder in ESM-2 SFT, and hyperparameters were learning rate=5e-5, batch size=8, and epoch=40. Spearman's correlation coefficient was used as a prediction accuracy measure. The bold letters indicate the better one from SVR and ESM-2 SFT.

| **Protein** | **function** | **Model** | **Exp only** | **ESM-2 zero-shot** | **Rosetta** | **Hybrid** |
|---|---|---|---|---|---|---|
| GRB2-SH3 | Binding | SVR | **0.5946** | **0.7195** | 0.6198 | 0.7182 |
| | | ESM-2 SFT | 0.5540 | 0.7186 | **0.6336** | **0.7388** |
| | Abundance | SVR | **0.6695** | **0.6551** | **0.6750** | **0.7170** |
| | | ESM-2 SFT | 0.5763 | 0.6405 | 0.6602 | 0.6910 |
| PSD95-PDZ | Binding | SVR | **0.6606** | **0.6520** | **0.6175** | **0.6775** |
| | | ESM-2 SFT | 0.5315 | 0.5914 | 0.5126 | 0.6191 |
| | Abundance | SVR | **0.6717** | **0.6675** | **0.6034** | **0.6541** |
| | | ESM-2 SFT | 0.4679 | 0.5511 | 0.5432 | 0.6138 |
| GB1 | Binding | SVR | 0.6485 | **0.6486** | **0.6633** | **0.6481** |
| | | ESM-2 SFT | **0.6646** | 0.3048 | 0.5002 | 0.435 |
| APH(3')II | Activity | SVR | **0.4185** | **0.5447** | **0.4998** | **0.5852** |
| | | ESM-2 SFT | 0.2840 | 0.5053 | 0.4250 | 0.5356 |
| CYP2C9 | Activity | SVR | **0.4577** | **0.6213** | **0.4346** | **0.6328** |
| | | ESM-2 SFT | 0.0990 | 0.1967 | 0.1056 | 0.1074 |
| DAOx | Activity | SVR | **0.3989** | **0.5719** | **0.4647** | **0.6266** |
| | | ESM-2 SFT | 0.1915 | 0.2219 | 0.2273 | 0.2926 |
| CcdB | Activity | SVR | **0.3749** | **0.4856** | **0.3873** | **0.4904** |
| | | ESM-2 SFT | 0.3470 | 0.4643 | 0.3640 | 0.4656 |
| KRAS | Abundance | SVR | **0.4881** | **0.4881** | **0.4881** | **0.4881** |
| | | ESM-2 SFT | 0.4111 | 0.1655 | 0.2543 | 0.2554 |
| FbFPs1 | Activity | SVR | **0.4403** | **0.6634** | **0.4553** | **0.6377** |
| | | ESM-2 SFT | 0.3351 | 0.6167 | 0.2970 | 0.6143 |
| DHFR | Activity | SVR | **0.3024** | **0.4629** | **0.3831** | 0.4724 |
| | | ESM-2 SFT | 0.2420 | 0.4977 | 0.2881 | **0.5006** |

Supplementary Table 3. The overlap between double-residue mutants in test data and their corresponding single-residue mutants in experimental training data. The position level overlap (Position) and the amino acid level overlap (Amino acid) were shown for each dataset and the amount of experimental training data. The results of GB1 were separately shown for models using descriptor and ESM-2 embedding since we used different test data for computational cost (Methods).

| **Protein** | **Functional value** | **Amount of experimental data** | **Position (%)** | **Amino acid (%)** |
|---|---|---|---|---|
| GRB2-SH3 | Binding affinity | 100 | 74.52 | 2.447 |
| | | 200 | 97.75 | 9.209 |
| | Abundance | 100 | 77.02 | 1.006 |
| | | 200 | 97.72 | 4.047 |
| PSD95-PDZ | Binding affinity | 100 | 51.67 | 0.357 |
| | | 200 | 85.64 | 1.863 |
| | Abundance | 100 | 45.15 | 0.901 |
| | | 200 | 84.71 | 2.191 |
| GB1(descriptor) | Binding affinity | 100 | 73.61 | 0.906 |
| | | 200 | 97.80 | 3.645 |
| GB1(ESM-2 embedding) | Binding affinity | 100 | 73.96 | 0.875 |
| | | 200 | 97.78 | 3.561 |
